# NetSolP: predicting protein solubility in E. coli using language models

**DOI:** 10.1101/2021.07.21.453084

**Authors:** Vineet Thumuluri, Hannah-Marie Martiny, Jose J. Almagro Armenteros, Jesper Salomon, Henrik Nielsen, Alexander R. Johansen

## Abstract

Solubility and expression levels of proteins can be a limiting factor for large-scale studies and industrial production. By determining the solubility and expression directly from the protein sequence, the success rate of wet-lab experiments can be increased.

In this study, we focus on predicting the solubility and usability for purification of proteins expressed in *Escherichia coli* directly from the sequence. Our model NetSolP is based on deep learning protein language models called transformers and we show that it achieves state-of-the-art performance and improves extrapolation across datasets. As we find current methods are built on biased datasets, we curate existing datasets by using strict sequence-identity partitioning and ensure that there is minimal bias in the sequences.

The predictor is available at https://services.healthtech.dtu.dk/service.php?NetSolP-1.0

## Introduction

Successful expression of soluble proteins is desired in research as well as commercial environments. High-throughput purification of proteins enables the production of various products in industries including pharmaceutical, food, and beverage [Chapman et al., 2018]. A large-scale protein structure determination effort^1^ has shown that low expression and solubility are common issues with about 49% successful expression rate for recombinant proteins and 52% purification rate for expressed proteins. There are several techniques that increase the solubility of wild-type proteins using mutations [Trevino et al., 2008, Miklos et al., 2012, Tan et al., 1998, Dudgeon et al., 2012, Costa et al., 2014]. Protein solubility depends on various external physical conditions such as pH and temperature and its interaction with intrinsic factors e.g., the amino-acid composition and structure of proteins. Reducing the search space to only protein sequences that potentially have high solubility and expression is beneficial to reduce the cost and time of wet-lab experiments. Thus, several sequence-based protein solubility prediction tools have been proposed using biophysical and structural features [Smialowski et al., 2012, Sormanni et al., 2015, Hebditch et al., 2017, Bhandari et al., 2020]. Recently, deep learning-based methods have been utilized to learn these features from the amino acid sequence only [Khurana et al., 2018, Raimondi et al., 2020, Wu and Yu, 2021]. Evolutionary data in the form of sequence profiles have proven valuable for producing high-quality predictors (Rawi et al., 2017, Khurana et al., 2018). However, computing Multiple Sequence Alignments (MSA) is slow and does not scale well for large numbers of proteins. All the above methods do not use the same objective. Camsol [Sormanni et al., 2015], Protein-Sol [Hebditch et al., 2017], SWI [Bhandari et al., 2020] predict the solubility, whereas, PROSO [Smialowski et al., 2012], DeepSol [Khurana et al., 2018], SKADE [Raimondi et al., 2020] and EPSOL [Wu and Yu, 2021] predict soluble expression of proteins.

Language models from Natural Language Processing have successfully been transferred to the protein domain due to the abundance of unlabelled raw sequence data. A protein language model, which is based on the transformer architecture [Vaswani et al., 2017], is trained in a self-supervised fashion on a large corpus, such as the UniRef50 database [Suzek et al., 2014], using the masked language-modelling objective [Devlin et al., 2019]. The transformer is a deep learning method to produce a contextual embedding of amino acids in the protein sequence. By using the masked language model objective it is able to build a context around each position and learns to “attend” or “focus” on amino acids and peptides that are relevant in the given context. These language models have been found to encode contact maps, taxonomy, and biophysical characteristics in their distributed representations [Rives et al., 2021, Rao et al., 2021, 2020, Elnaggar et al., 2020, Vig et al., 2020, Brandes et al., 2021, Martiny et al., 2021]. In this study, we use a protein language model to predict two objectives, solubility and practical usability for purification of proteins in *E. coli*, and obtain state-of-the-art performance. As we find current datasets are biased by artifacts introduced by the expression vector, we also curate multiple protein datasets for both objectives from publicly available data. Our curation, using strict homology partitioning and ensuring no sequence bias, makes them a better representative of real-world performance than current datasets.

## Data

### TargetTrack dataset

Rawi et al. [2017] curated 69,420 proteins as the training set from a larger collection of 129,643 proteins (Smialowski et al., 2012) and used 2001 protein sequences curated by Chang et al. [2013] as an independent test set. All of these sequences were selected from the TargetTrack database (Berman et al., 2017), which was a large-scale project by the Protein Structure Initiative (PSI) from 2000-2017 to greatly increase the number of known protein structures. No explicit solubility label is recorded in the database, although several participating centres registered it separately [Seiler et al., 2014] and thus the binary solubility label for some proteins is available in sources such as the PSI: Biology dataset described below. Proteins from the downloaded version were considered soluble by Smialowski et al. [2012] if they reached a set of predetermined soluble experimental states and insoluble if they did not reach those states in the version released 8 months later and also did not already have a structure submitted to the Protein Data Bank (PDB).

### Biases in the TargetTrack dataset

The PaRSnIP (Rawi et al., 2017), DeepSol (Khurana et al., 2018), and SKADE (Raimondi et al., 2020) soluble expression predictors were built using the curated train set and were shown to achieve high scores on the test set. However, it was noticed that these tools generalize poorly (Bhandari et al., 2020, Hon et al., 2021). Raimondi et al. [2020] showed that the SKADE model focused mostly on the N- and C- termini and validated that DeepSol did the same using an experiment that involved cropping the starting and ending segments of the sequences. Unfortunately, this behaviour is likely not due to underlying biophysics but a result of unintended bias in both the training and test sets. An example of this is that 11,602 out of 69,420 sequences of the training set and 344 out of 2001 sequences of the test set have the N-terminal His-tag ‘MGSDKIHHHHHH’ with ~99% and ~97% of them being insoluble respectively. His-tags are polyhistidine peptides incorporated in the recombinant protein to enable affinity purification [Spriestersbach et al., 2015]. Other N-terminal His-tags like ‘MGSSHHHH’, ‘MHHHHHHS’, ‘MRGSHHHH’ with over 100 instances each have 88%, 100%, and 100% mean solubility, respectively. An example of a C-terminal His-tag from the dataset is ‘HHHHH’, when sequences have an amino acid other than E preceding this His-tag, they are almost always soluble. Such statistics are not expected naturally and indicate a bias in the selection or the experiments themselves. Moreover, Hon et al. [2021] compared the labels of sequences from this dataset with another dataset whose solubility was provided separately (Price et al., 2011) and found that around 18.6% of labels were different, even with 100% identical sequences. The consequence of this is that the trained models focus more on the His-tag instead of the wild-type sequence. Since the SoluProt and EPSOL [Wu and Yu, 2021] training datasets also come from the same source, they face the same issues discussed above. Therefore, the biased selection of proteins in combination with the label noise makes it difficult to train generalizable models using this dataset.

### PSI: Biology dataset

As part of PSI, several centres recorded explicit expression and solubility labels for the target proteins. A subset of this data was extracted by Bhandari et al. [2020], which had 12,216 proteins expressed in *E. coli* using two specific expression vectors ‘pET21’ and ‘pET15’. Although newer techniques can change the status of some proteins, explicit labels make this dataset far more reliable. The average solubility of the proteins is ~66%. We use this dataset for 5-fold cross-validation.

### Price dataset

The North East Structural Consortium (NESG) expressed 9644 proteins in *E. coli* using a unified production pipeline (Price et al., 2011) and provide integer scores (0-5) for both expression (E) and solubility (S). The proteins are part of the TargetTrack database, but the scores were obtained by Hon et al. [2021] from the original authors. We remove sequences that have multiple scores and use the remaining 9272 sequences in two ways. First, as an independent test set for solubility consisting of 1323 highly expressed proteins (E = 4 or 5) with high solubility score (S = 4 or 5) as soluble and low score (S = 0) as insoluble. Using this definition, soluble proteins are ~64% of the test set. An alternative objective ‘usability’, which requires the protein to be successfully purified on a large scale, is used to generate a new dataset. Usability is estimated using the product U=E.S by the authors. Proteins are considered usable if U is greater than 11 and unusable if U is less than 4. The total number of proteins for 5-fold cross-validation is 7,259. We exclude proteins with intermediate scores in both solubility and usability datasets to reduce potential noise.

### Camsol mutation dataset

Sormanni et al., 2015 compiled a set of 19 proteins with 56 total variants from four sources whose change in solubility was experimentally verified. Compared to the wild-type, 53 mutations increased solubility and 3 decreased it. This dataset is used as an independent test set and no partitioning is performed.

## Methods

### Data Partitioning

To generate high-quality data partitions, we use the four-phase procedure described in Gíslason et al., 2021 to make label-balanced splits for 5-fold cross-validation. This procedure ensures that each pair of train and test fold does not share sequences that have global sequence identity greater than 25% as determined using ggsearch36, which is a part of the FASTA package (Pearson and Lipman, 1988). The datasets after partitioning are as follows, PSI: Biology solubility cross-validation set with 11,226 sequences, the Price usability cross-validation set with 7,259 sequences, and the Price solubility independent test set with 1,323 sequences. The latter dataset is ensured not to share sequences with global identity greater than 25% with the full PSI: Biology dataset using USEARCH v11.0.667, 32-bit (Edgar, 2010).

### NetSolP

Multiple publicly available transformer models are evaluated. We refer to the 12-layer ESM (Evolutionary Scale Modelling, Rives et al., 2021) model with 84M parameters as ESM12, the 12-layer ESM model using multiple sequence alignments (Rao et al., 2021) with 100M parameters as ESM-MSA, the 33-layer ESM model with 650M parameters as ESM1b (Rao et al., 2020) and the 24-layer ProtT5-XL-UniRef50 encoder model (Elnaggar et al., 2020) with 1208M parameters as ProtT5. We follow the guidelines of [Rao et al., 2021] for generating MSAs. For each protein sequence, we construct an MSA using HHblits, version 3.1.0. ([Steinegger et al., 2019]) against the UniClust30_2020−6_ database ([Mirdita et al., 2016]) with default settings except setting number of iterations to 3 (−*n* 3). To reduce the size of MSA and memory requirements, hhfilter is applied ([Steinegger et al., 2019]) with the -diff 64 parameter. The original MSA transformers uses 256.

The output representations of each amino acid in the sequence are averaged to represent the protein and a linear classification layer is used to predict binary solubility. The trained models have a suffix ‘−F’ and ‘−P’ to indicate whether they are trained end-to-end (fine-tuning) or only the classification layer (pretrained embedding), which is based on the available computational resources. The maximum sequence length used for training is 510, by removing around 3.4% of the training sequences that exceed this length, to speed up the training process. For prediction, amino acids after position 1022 are removed due to the maximum length constraints of the transformer models. Different learning rates for the transformer (3×10^−6^) and classification layer (2×10^−5^) are used, and the training is terminated using early stopping. Mixed-precision and model sharding techniques are utilized to efficiently fine-tune the models. The PyTorch-lightning (Falcon et al., 2019) library and hardware provided by Google Colaboratory GPUs^2^, and 2 Tesla V100s are used for training and testing. We improve the speed and memory utilization of the final tool using ONNX-runtime^3^ and dynamic quantization. The final predictor (NetSolP) is an ensemble of fine-tuned, dynamically quantized ESM1b models. Additionally, we provide a distilled [Hinton et al., 2015] version, NetSolP-D, that preserves most of the performance but runs five times as fast.

For qualitative analysis, we calculate the contributions of amino acids in the sequence towards predictions, for the ESM12 ensemble, using Integrated Gradients method (Sundararajan et al., 2017) from the Captum^4^ library. The baseline is taken to be a < *CLS* > token followed by < *PAD* > tokens i.e. an empty protein sequence. Other parameters are set to their default values. The per amino acid contribution for each model in the ensemble is summed and then normalized over the protein sequence using the L1-Norm. The importance is taken to be the absolute value of the contributions. For calculating conserved residue scores the tool provided by Capra and Singh [2007]^5^ is used with scaled Shannon entropy and a window-size 0. The protein families are chosen from the PSI: Biology training set using MMseqs2 (Steinegger and Söding, 2017) with a minimum sequence identity 0.2 and the coverage set to 0.5. Three families, FAD/NAD(P)-binding domain (InterPro domain IPR036188), DNA breaking-rejoining enzyme, catalytic core (InterPro domain IPR011010) and Trehalase (Panther domain PTHR31616), are selected such that they have many sequences (48, 71, and 50 respectively) and have average solubility close to 50% (56%, 51% and, 41% respectively).

## Results & Discussion

We compare PaRSnIP, Camsol, DeepSol-S2, ProteinSol, SWI, SoluProt, and multiple transformer models using threshold-dependent metrics such as accuracy, precision, Matthew’s correlation coefficient (MCC), and a threshold-independent metric, area under the ROC curve (AUC). For most models the value recommended by the authors is used as the threshold for predicting soluble proteins. Since Camsol is not built for binary predictions we use a value of 1. The threshold used in cross-validation for each model of the NetSolP ensemble is 0.5 and the threshold for NetSolP is set as the average of optimal thresholds for each of the five validation folds, computed using the Youden Index (Youden, 1950). PaRSnIP, DeepSol-S2, and ESM-MSA require sequence profiles as input and thus are infeasible for predicting a large set of proteins. Camsol, ProteinSol, and SWI require only the protein sequence and thus scale well. SWI (Bhandari et al., 2020) and SoluProt (Hon et al., 2021) are retrained to better represent the scores with our modified dataset splits.

NetSolP outperforms existing solubility prediction tools on the PSI: Biology “Solubility” 5-fold cross-validation dataset (Table 1) of 11,226 sequences, with the highest AUC (0.73 ± 0.02), MCC (0.29 ± 0.04) and accuracy (0.70 ± 0.02). Among transformer models the best scores are obtained by ESM-MSA which uses sequence profiles. The independent validation set (Table 2), compared among predictors that do not use sequence profiles, shows that NetSolP also generalizes better with the highest AUC (0.760), MCC (0.402), accuracy (0.728) and precision (0.773). NetSolP-D, which is the distilled version of the NetSolP ensemble, performs almost as well with AUC (0.756), MCC (0.391), accuracy (0.723) and precision (0.769)

**Table 1.** PSI:Biology Solubility. 5-Fold CV

| Models | ACC | PRE | MCC | AUC |
| --- | --- | --- | --- | --- |
| SoluProt | 0.59 ± 0.03 | 0.70 ± 0.02 | 0.10 ± 0.03 | 0.59 ± 0.02 |
| Parsnip* | 0.61 ± 0.10 | 0.71 ± 0.03 | 0.16 ± 0.11 | 0.64 ± 0.08 |
| Camsol | 0.59 ± 0.08 | 0.77 ± 0.03 | 0.21 ± 0.09 | 0.65 ± 0.06 |
| DeepSol S2* | 0.54 ± 0.02 | **0.81 ± 0.05** | 0.22 ± 0.05 | 0.67 ± 0.04 |
| ProteinSol | 0.70 ± 0.03 | 0.70 ± 0.03 | 0.23 ± 0.08 | 0.68 ± 0.05 |
| SWI | 0.64 ± 0.04 | 0.80 ± 0.02 | 0.29 ± 0.05 | 0.69 ± 0.04 |
| SWI† | 0.63 ± 0.04 | 0.79 ± 0.03 | 0.28 ± 0.05 | 0.69 ± 0.03 |
| ESM12-F | **0.71 ± 0.03** | 0.75 ± 0.03 | 0.32 ± 0.04 | 0.73 ± 0.04 |
| ESM1b-F | 0.70 ± 0.02 | 0.74 ± 0.03 | 0.29 ± 0.03 | 0.73 ± 0.03 |
| ProtT5-P | 0.70 ± 0.03 | 0.77 ± 0.02 | **0.33 ± 0.04** | 0.73 ± 0.02 |
| ESM-MSA-P* | **0.71 ± 0.03** | 0.76 ± 0.02 | **0.33 ± 0.04** | **0.75 ± 0.03** |
| NetSolP | 0.70 ± 0.02 | 0.74 ± 0.03 | 0.29 ± 0.04 | 0.73 ± 0.02 |

†= Retrained on this dataset
\* = Method requires sequence profiles

**Table 2.** Price Solubility. Independent Validation

| Method | ACC | PRE | MCC | AUC |
| --- | --- | --- | --- | --- |
| SoluProt | 0.624 | 0.704 | 0.187 | 0.634 |
| Camsol | 0.570 | 0.751 | 0.199 | 0.646 |
| Protein-Sol | 0.641 | 0.694 | 0.190 | 0.679 |
| SWI | 0.680 | 0.712 | 0.269 | 0.690 |
| ESM12-F | 0.698 | 0.743 | 0.328 | 0.732 |
| ESM1b-F | 0.723 | 0.764 | 0.386 | **0.761** |
| ProtT5-P | 0.702 | 0.714 | 0.314 | 0.733 |
| ESM-MSA-P | 0.712 | 0.715 | 0.340 | 0.745 |
| NetSolP | **0.728** | **0.773** | **0.402** | 0.760 |
| NetSolP-D | 0.723 | 0.769 | 0.391 | 0.756 |

Interestingly, NetSolP is unable to discriminate between the minute solubility changes produced by mutations compared to other methods (Table 3). However, a high accuracy (94.6%) by ESM12 shows that it may be more suitable for comparing highly similar proteins. Only SODA (Paladin et al., 2017) was trained with the goal of predicting solubility changes upon point mutations, unlike the rest which used the binary solubility objective.

**Table 3.** Camsol Solubility Mutation. Independent Validation

| Method | Trevino | Miklos | Tan | Dudgeon | Total | Accuracy |
| --- | --- | --- | --- | --- | --- | --- |
| SoluProt | 17 / 22 | 3 / 3 | 1 / 1 | 11 / 30 | 32 / 56 | 57.1 |
| PROSO II | 16 / 22 | 3 / 3 | 1 / 1 | 12 / 30 | 32 / 56 | 57.1 |
| SolPro | 15 / 22 | 3 / 3 | 1 / 1 | 21 / 30 | 40 / 56 | 71.4 |
| CamSol | 22 / 22 | 3 / 3 | 1 / 1 | 28 / 30 | 54 / 56 | 96.4 |
| SWI | 21 / 22 | 3 / 3 | 1 / 1 | 30 / 30 | 55 / 56 | 98.2 |
| SODA | 22 / 22 | 3 / 3 | 1 / 1 | 30 / 30 | 56 / 56 | 100.0 |
| ESM12-F | 19 / 22 | 3 / 3 | 1 / 1 | 30 / 30 | 53 / 56 | 94.6 |
| ESM1b-F | 14 / 22 | 3 / 3 | 0 / 1 | 14 / 30 | 31 / 56 | 55.3 |
| NetSolP | 16 / 22 | 3 / 3 | 0 / 1 | 18 / 30 | 37 / 56 | 66.1 |

On the Price et al. [2011] dataset (Table 4) with the “Usability” objective the highest AUC (0.71 ± 0.01), MCC (0.30 ± 0.03), precision (0.64 ± 0.03) and accuracy (0.65 ± 0.01) is obtained by NetSolP. Quantization proves to be very effective as NetSolP is able to retain most of the performance of the constituent ESM1b models while reducing its data storage by a factor of four.

**Table 4.** Price Usability. 5-Fold CV

| Method | ACC | PRE | MCC | AUC |
| --- | --- | --- | --- | --- |
| SoluProt† | 0.63 ± 0.02 | 0.63 ± 0.03 | 0.26 ± 0.03 | 0.63 ± 0.02 |
| SWI | 0.58 ± 0.01 | 0.54 ± 0.01 | 0.20 ± 0.02 | 0.64 ± 0.02 |
| SoluProt | 0.62 ± 0.01 | 0.60 ± 0.02 | 0.24 ± 0.02 | 0.67 ± 0.02 |
| ESM12-F | 0.64 ± 0.01 | 0.63 ± 0.01 | 0.29 ± 0.03 | **0.71 ± 0.01** |
| ESM1b-F | **0.65 ± 0.01** | **0.64 ± 0.03** | **0.30 ± 0.03** | **0.71 ± 0.01** |
| NetSolP | 0.65 ± 0.02 | 0.65 ± 0.04 | **0.30 ± 0.04** | 0.70 ± 0.01 |

†= Retrained on this dataset

Fig 1 shows that the solubility of shorter sequences is predicted slightly better than that of longer sequences by our method as well as by SWI which could be due to the abundance of shorter sequences in the datasets. Raimondi et al., 2020 observe that the ends of the protein sequence are more important for predicting the solubility. However, in our case (Fig 2) the magnitude of the effect is insignificant, with the exception of the initial amino acid. The initial 1% of the amino acid sequence has 3% of the total importance indicating that it is only a small bias. The signed contributions averaged over all the positions for an amino acid show an interesting relationship between NetSolP and SWI. The spearman rank correlation for these two sets of amino acid solubility scores is 0.66 (p-value= 1.48 × 10^−3^) suggesting that the average statistics learned are similar but the performance improvement for NetSolP over SWI could be due to the context-dependent contribution of amino acids towards solubility. The conservation versus importance plots (Fig 3) show that highly conserved regions tend to be more important but not vice versa.

**Figure 1.**
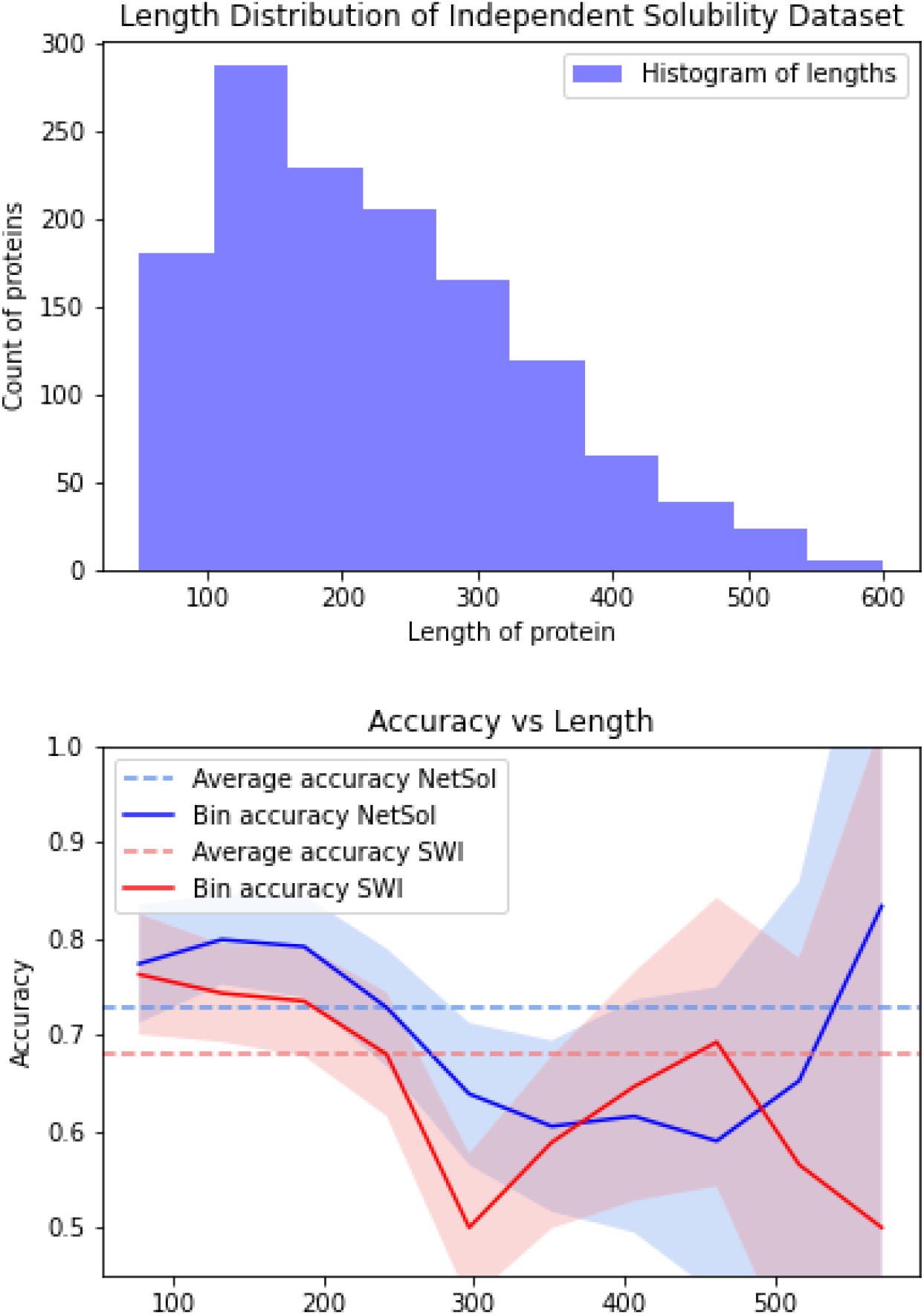
*Top*: The length distribution of the test set. *Bottom*: Change in accuracy based on the length of the protein sequences computed on the Price Solubility independent validation set of 1323 sequences. A dip in accuracy with longer sequences can be seen for both NetSolP and the best existing tool SWI

**Figure 2.**
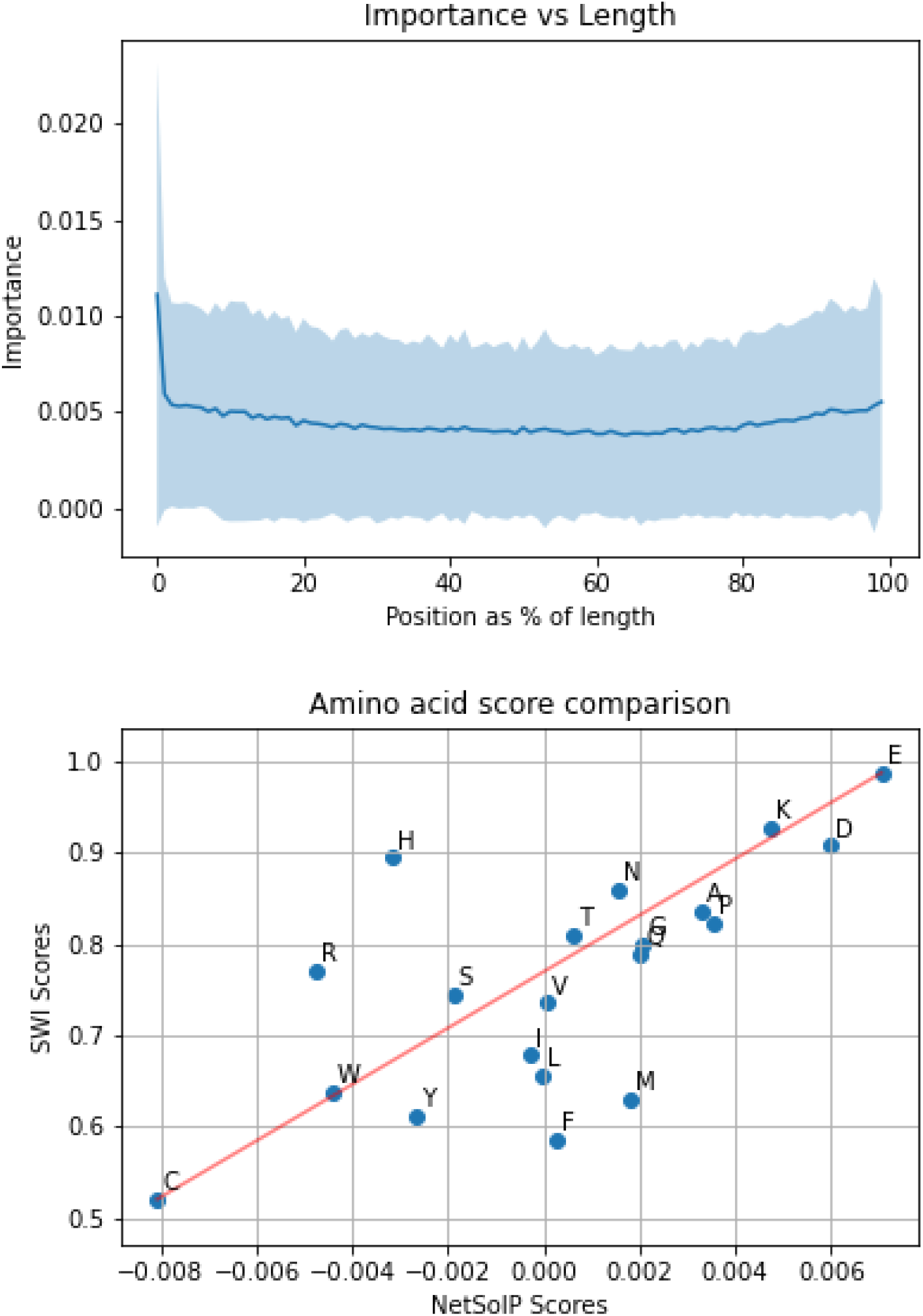
Qualitative analysis of ESM12 model using Integrated Gradients computed on the independent solubility dataset. *Top*: The importance of position in solubility prediction. *Bottom*: Solubility scores per amino acid shows strong correlation between NetSolP and SWI.

**Figure 3.**
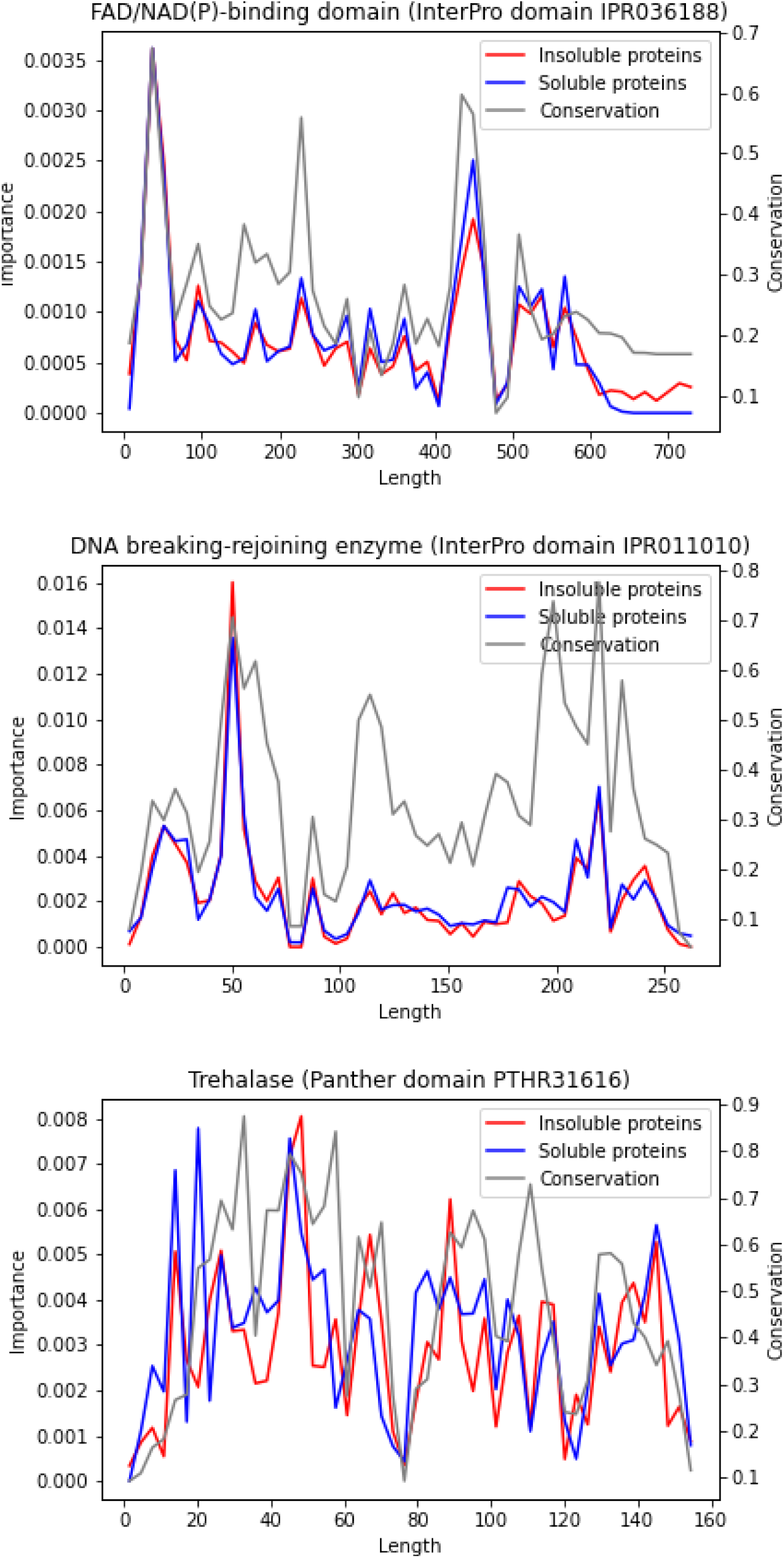
Plotting the residue conservation score and importance for each position in the aligned protein families shows that highly conserved regions are important but not vice versa. Three protein families from the training set were chosen.

## Conclusion

We propose NetSolP, a predictor based on protein language models and deep learning, that outperforms existing tools for *in silico* solubility and usability prediction. We curate new datasets with an emphasis on strict partitioning based on sequence identity and ensuring that there are no spurious correlations between the sequences and target labels. Our experiments find that larger transformer models are better and fine-tuning the models is more effective than the pretrained embeddings. Qualitative analysis reveals an interesting correlation with a previous method SWI and finds no significantly important regions for solubility in contrast to the study done by Raimondi et al., 2020, which can be explained by the bias present in the dataset they use. The predictor is available at https://services.healthtech.dtu.dk/service.php?NetSolP-1.0.

## Footnotes

1 http://targetdb.rcsb.org/metrics

2 https://colab.research.google.com/

3 https://github.com/microsoft/onnxruntime

4 https://captum.ai/

5 https://compbio.cs.princeton.edu/conservation/score.html

## References

Jordan M Chapman, A. E. Ismail, and C. Dinu. Industrial applications of enzymes: Recent advances, techniques, and outlooks. Catalysts, 8:238, 2018.

Saul R. Trevino, J.Martin Scholtz, and C.Nick Pace. Measuring and Increasing Protein Solubility. Journal of Pharmaceutical Sciences, 97(10):4155–4166, October 2008. ISSN 00223549. doi: 10.1002/jps.21327. URL https://linkinghub.elsevier.com/retrieve/pii/S0022354916327514.

Aleksandr E. Miklos, Christien Kluwe, Bryan S. Der, Supriya Pai, Aroop Sircar, Randall A. Hughes, Monica Berrondo, Jianqing Xu, Vlad Codrea, Patricia E. Buckley, Alena M. Calm, Heather S. Welsh, Candice R. Warner, Melody A. Zacharko, James P. Carney, Jeffrey J. Gray, George Georgiou, Brian Kuhlman, and Andrew D. Ellington. Structure-based design of supercharged, highly thermoresistant antibodies. Chemistry & Biology, 19(4):449–455, 2012. ISSN 1074-5521. doi: https://doi.org/10.1016/j.chembiol.2012.01.018. URL https://www.sciencedirect.com/science/article/pii/S1074552112000774.

Philip H. Tan, Vano Chu, James E. Stray, Donald K. Hamlin, Dean Pettit, D.Scott Wilbur, Robert L. Vessella, and Patrick S. Stayton. Engineering the isoelectric point of a renal cell carcinoma targeting antibody greatly enhances scfv solubility. Immunotechnology, 4(2):107–114, 1998. ISSN 1380-2933. doi: https://doi.org/10.1016/S1380-2933(98)00011-6. URL https://www.sciencedirect.com/science/article/pii/S1380293398000116.

Kip Dudgeon, Romain Rouet, Iris Kokmeijer, Peter Schofield, Jessica Stolp, David Langley, Daniela Stock, and Daniel Christ. General strategy for the generation of human antibody variable domains with increased aggregation resistance. Proceedings of the National Academy of Sciences, 109(27):10879–10884, 2012. ISSN 0027-8424. doi: 10.1073/pnas.1202866109. URL https://www.pnas.org/content/109/27/10879.

Sofia Costa, André Almeida, António Castro, and Lucília Domingues. Fusion tags for protein solubility, purification and immunogenicity in Escherichia coli: the novel Fh8 system. Frontiers in Microbiology, 5, 2014. ISSN 1664-302X. doi: 10.3389/fmicb.2014.00063. URL http://journal.frontiersin.org/article/10.3389/fmicb.2014.00063/abstract.

Pawel Smialowski, Gero Doose, Phillipp Torkler, Stefanie Kaufmann, and Dmitrij Frishman. PROSO II - a new method for protein solubility prediction. FEBS Journal, 279(12):2192–2200, May 2012. doi: 10.1111/j.1742-4658.2012.08603.x. URL http://dx.doi.org/10.1111/j.1742-4658.2012.08603.x.

Pietro Sormanni, Francesco A. Aprile, and Michele Vendruscolo. The camsol method of rational design of protein mutants with enhanced solubility. Journal of Molecular Biology, 427(2):478–490, 2015. ISSN 0022-2836. doi: https://doi.org/10.1016/j.jmb.2014.09.026. URL https://www.sciencedirect.com/science/article/pii/S0022283614005312.

Max Hebditch, M Alejandro Carballo-Amador, Spyros Charonis, Robin Curtis, and Jim Warwicker. Protein–Sol: a web tool for predicting protein solubility from sequence. Bioinformatics, 33(19):3098–3100, 05 2017. ISSN 1367-4803. doi: 10.1093/bioinformatics/btx345. URL https://doi.org/10.1093/bioinformatics/btx345.

Bikash K Bhandari, Paul P Gardner, and Chun Shen Lim. Solubility-Weighted Index: fast and accurate prediction of protein solubility. Bioinformatics, 36(18):4691–4698, 06 2020. ISSN 1367-4803. doi: 10.1093/bioinformatics/btaa578. URL https://doi.org/10.1093/bioinformatics/btaa578.

Sameer Khurana, Reda Rawi, Khalid Kunji, Gwo-Yu Chuang, Halima Bensmail, and Raghvendra Mall. DeepSol: a deep learning framework for sequence-based protein solubility prediction. Bioinformatics, 34(15):2605–2613, 03 2018. doi: 10.1093/bioinformatics/bty166. URL https://doi.org/10.1093/bioinformatics/bty166.

Daniele Raimondi, Gabriele Orlando, Piero Fariselli, and Yves Moreau. Insight into the protein solubility driving forces with neural attention. PLOS Computational Biology, 16(4):1–15, 04 2020. doi: 10.1371/journal.pcbi.1007722. URL https://doi.org/10.1371/journal.pcbi.1007722.

Xiang Wu and Liang Yu. EPSOL: sequence-based protein solubility prediction using multidimensional embedding. Bioinformatics, 06 2021. ISSN 1367-4803. doi: 10.1093/bioinformatics/btab463. URL https://doi.org/10.1093/bioinformatics/btab463.btab463.

Reda Rawi, Raghvendra Mall, Khalid Kunji, Chen-Hsiang Shen, Peter D Kwong, and Gwo-Yu Chuang. PaRSnIP: sequence-based protein solubility prediction using gradient boosting machine. Bioinformatics, 34(7):1092–1098, 10 2017. ISSN 1367-4803. doi: 10.1093/bioinformatics/btx662. URL https://doi.org/10.1093/bioinformatics/btx662.

Ashish Vaswani, Noam Shazeer, Niki Parmar, Jakob Uszkoreit, Llion Jones, Aidan N. Gomez, Lukasz Kaiser, and Illia Polosukhin. Attention is all you need, 2017.

Baris E. Suzek, Yuqi Wang, Hongzhan Huang, Peter B. McGarvey, Cathy H. Wu, and the UniProt Consortium. UniRef clusters: a comprehensive and scalable alternative for improving sequence similarity searches. Bioinformatics, 31(6):926–932, 11 2014. ISSN 1367-4803. doi: 10.1093/bioinformatics/btu739. URL https://doi.org/10.1093/bioinformatics/btu739.

Jacob Devlin, Ming-Wei Chang, Kenton Lee, and Kristina Toutanova. Bert: Pre-training of deep bidirectional transformers for language understanding, 2019.

Alexander Rives, Joshua Meier, Tom Sercu, Siddharth Goyal, Zeming Lin, Jason Liu, Demi Guo, Myle Ott, C. Lawrence Zitnick, Jerry Ma, and Rob Fergus. Biological structure and function emerge from scaling unsupervised learning to 250 million protein sequences. Proceedings of the National Academy of Sciences, 118(15): e2016239118, 2021. doi: 10.1073/pnas.2016239118.

Roshan Rao, Jason Liu, Robert Verkuil, Joshua Meier, John F. Canny, Pieter Abbeel, Tom Sercu, and Alexander Rives. Msa transformer. bioRxiv, page 2021.02.12.430858, 2021. doi: 10.1101/2021.02.12.430858. URL https://www.biorxiv.org/content/10.1101/2021.02.12.430858v1.

Roshan Rao, Joshua Meier, Tom Sercu, Sergey Ovchinnikov, and Alexander Rives. Transformer protein language models are unsupervised structure learners. bioRxiv, page 2020.12.15.422761, 2020. doi: 10.1101/2020.12.15.422761. URL https://www.biorxiv.org/content/10.1101/2020.12.15.422761v1.

Ahmed Elnaggar, Michael Heinzinger, Christian Dallago, Ghalia Rehawi, Yu Wang, Llion Jones, Tom Gibbs, Tamas Feher, Christoph Angerer, Martin Steinegger, DEBSINDHU Bhowmik, and Burkhard Rost. Prottrans: Towards cracking the language of life’s code through self-supervised deep learning and high performance computing. bioRxiv, page 2020.07.12.199554, 2020. doi: 10.1101/2020.07.12.199554. URL https://www.biorxiv.org/content/early/2020/07/21/2020.07.12.199554.

Jesse Vig, Ali Madani, Lav R. Varshney, Caiming Xiong, Richard Socher, and Nazneen Fatema Rajani. Bertology meets biology: Interpreting attention in protein language models. arXiv, page 2006.15222, 2020. URL https://arxiv.org/abs/2006.15222.

Nadav Brandes, Dan Ofer, Yam Peleg, Nadav Rappoport, and Michal Linial. Proteinbert: A universal deep-learning model of protein sequence and function. bioRxiv, 2021. doi: 10.1101/2021.05.24.445464. URL https://www.biorxiv.org/content/early/2021/05/25/2021.05.24.445464.

Hannah-Marie Martiny, Jose Juan Almagro Armenteros, Alexander Rosenberg Johansen, Jesper Salomon, and Henrik Nielsen. Deep protein representations enable recombinant protein expression prediction. bioRxiv, 2021. doi: 10.1101/2021.05.13.443426. URL https://www.biorxiv.org/content/early/2021/05/14/2021.05.13.443426.

Catherine Ching Han Chang, Jiangning Song, Beng Ti Tey, and Ramakrishnan Nagasundara Ramanan. Bioinformatics approaches for improved recombinant protein production in Escherichia coli: protein solubility prediction. Briefings in Bioinformatics, 15(6):953–962, 08 2013. ISSN 1467-5463. doi: 10.1093/bib/bbt057. URL https://doi.org/10.1093/bib/bbt057.

Helen M. Berman, Margaret J. Gabanyi, and Protein Structure Initiative Network Of Investigators. Protein Structure Initiative - Targettrack 2000-2017 - All Data Files, July 2017. URL https://zenodo.org/record/821654. https://zenodo.org/record/821654.

Catherine Y. Seiler, Jin G. Park, Amit Sharma, Preston Hunter, Padmini Surapaneni, Casey Sedillo, James Field, Rhys Algar, Andrea Price, Jason Steel, Andrea Throop, Michael Fiacco, and Joshua LaBaer. DNASU plasmid and PSI:Biology-Materials repositories: resources to accelerate biological research. Nucleic Acids Research, 42 (D1):D1253–D1260, January 2014. ISSN 0305-1048, 1362-4962. doi: 10.1093/nar/gkt1060. URL https://academic.oup.com/nar/article-lookup/doi/10.1093/nar/gkt1060.

Jiri Hon, Martin Marusiak, Tomas Martinek, Antonin Kunka, Jaroslav Zendulka, David Bednar, and Jiri Damborsky. SoluProt: prediction of soluble protein expression in Escherichia coli. Bioinformatics, 37(1):23–28, 01 2021. ISSN 1367-4803. doi: 10.1093/bioinformatics/btaa1102. URL https://doi.org/10.1093/bioinformatics/btaa1102.

Anne Spriestersbach, Jan Kubicek, Frank Schäfer, Helena Block, and Barbara Maertens. Chapter one - purification of his-tagged proteins. In Jon R. Lorsch, editor, Laboratory Methods in Enzymology: Protein Part D, volume 559 of Methods in Enzymology, pages 1–15. Academic Press, 2015. doi: https://doi.org/10.1016/bs.mie.2014.11.003. URL https://www.sciencedirect.com/science/article/pii/S0076687914000688.

W Nicholson Price, Samuel K Handelman, John K Everett, Saichiu N Tong, Ana Bracic, Jon D Luff, Victor Naumov, Thomas Acton, Philip Manor, Rong Xiao, Burkhard Rost, Gaetano T Montelione, and John F Hunt. Large-scale experimental studies show unexpected amino acid effects on protein expression and solubility in vivo in E. coli. Microbial Informatics and Experimentation, 1(1):6, December 2011. ISSN 2042-5783. doi: 10.1186/2042-5783-1-6. URL https://microbialinformaticsj.biomedcentral.com/articles/10.1186/2042-5783-1-6.

Magnús Halldór Gíslason, Henrik Nielsen, José Juan Almagro Armenteros, and Alexander Rosenberg Johansen. Prediction of gpi-anchored proteins with pointer neural networks. Current Research in Biotechnology, 3:6–13, 2021. ISSN 2590-2628. doi: https://doi.org/10.1016/j.crbiot.2021.01.001. URL https://www.sciencedirect.com/science/article/pii/S2590262821000010.

W. R. Pearson and D. J. Lipman. Improved tools for biological sequence comparision. Proc. Natl. Acad. Sci., 85:2444–2448, 1988.

Robert C. Edgar. Search and clustering orders of magnitude faster than BLAST. Bioinformatics, 26(19):2460–2461, 08 2010. ISSN 1367-4803. doi: 10.1093/bioinformatics/btq461. URL https://doi.org/10.1093/bioinformatics/btq461.

Martin Steinegger, Markus Meier, Milot Mirdita, Harald Vöhringer, Stephan J. Haunsberger, and Johannes Söding. Hh-suite3 for fast remote homology detection and deep protein annotation. bioRxiv, 2019. doi: 10.1101/560029. URL https://www.biorxiv.org/content/early/2019/02/25/560029.

Milot Mirdita, Lars von den Driesch, Clovis Galiez, Maria J. Martin, Johannes Söding, and Martin Steinegger. Uniclust databases of clustered and deeply annotated protein sequences and alignments. Nucleic Acids Research, 45(D1):D170–D176, 11 2016. ISSN 0305-1048. doi: 10.1093/nar/gkw1081. URL https://doi.org/10.1093/nar/gkw1081.

WA Falcon et al. Pytorch lightning. GitHub. Note: https://github.com/PyTorchLightning/pytorch-lightning, 3, 2019.

Geoffrey Hinton, Oriol Vinyals, and Jeff Dean. Distilling the knowledge in a neural network, 2015.

Mukund Sundararajan, Ankur Taly, and Qiqi Yan. Axiomatic attribution for deep networks. arXiv, page 1703.01365, 2017. URL http://arxiv.org/abs/1703.01365.

John A. Capra and Mona Singh. Predicting functionally important residues from sequence conservation. Bioinformatics, 23(15):1875–1882, 05 2007. ISSN 1367-4803. doi: 10.1093/bioinformatics/btm270. URL https://doi.org/10.1093/bioinformatics/btm270.

Martin Steinegger and Johannes Söding. MMseqs2 enables sensitive protein sequence searching for the analysis of massive data sets. Nature Biotechnology, 35(11): 1026–1028, November 2017. ISSN 1087-0156, 1546-1696. doi: 10.1038/nbt.3988. URL http://www.nature.com/articles/nbt.3988.

W. J. Youden. Index for rating diagnostic tests. Cancer, 3(1):32–35, 1950. doi: https://doi.org/10.1002/1097-0142(1950)3:1?32::AID-CNCR2820030106?3.0.CO;2-3. URL https://acsjournals.onlinelibrary.wiley.com/doi/abs/10.1002/1097-0142%281950%293%3A1%3C32%3A%3AAID-CNCR2820030106%3E3.0.CO%3B2-3.

Lisanna Paladin, Damiano Piovesan, and Silvio C. E. Tosatto. SODA: prediction of protein solubility from disorder and aggregation propensity. Nucleic Acids Research, 45(W1):W236–W240, 05 2017. ISSN 0305-1048. doi: 10.1093/nar/gkx412. URL https://doi.org/10.1093/nar/gkx412.

